# Corrective muscle activity reveals subject-specific sensorimotor recalibration

**DOI:** 10.1101/372359

**Authors:** Pablo A. Iturralde, Gelsy Torres-Oviedo

## Abstract

Recent studies suggest that planned and corrective actions are recalibrated during some forms of motor adaptation. However, corrective (a.k.a., reactive) movements in human locomotion are thought to simply reflect sudden environmental changes independently from sensorimotor recalibration. Thus, we asked if corrective responses can indicate the motor system’s adapted state following prolonged exposure to a novel walking situation inducing sensorimotor adaptation. We recorded electromyographic signals bilaterally on 15 leg muscles before, during, and after split-belts walking (i.e., novel walking situation), in which the legs move at different speeds. We exploited the rapid temporal dynamics of corrective responses upon unexpected speed transitions to isolate them from the overall motor output. We found that corrective muscle activity was structurally different following short vs. long exposures to split-belts walking. Only after a long exposure, removal of the novel environment elicited corrective muscle patterns that matched those expected in response to a perturbation opposite to the one originally experienced. This indicated that individuals who recalibrated their motor system adopted split-belts environment as their new “normal” and transitioning back to the original walking environment causes subjects to react as if it was novel to them. Interestingly, this learning declined with age, but steady state modulation of muscle activity during split-belts walking did not, suggesting potentially different neural mechanisms underlying these motor patterns. Taken together, our results show that corrective motor commands reflect the adapted state of the motor system, which is less flexible as we age.

**Significance statement:** We showed that corrective muscle activity elicited by sudden environmental transitions are revealing of the underlying recalibration process during sensorimotor adaptation. Our finding provides an alternate metric for quantifying the motor system’s recalibration other than aftereffects or post-adaptation errors, which might not be clearly defined in some tasks. Notably, our novel approach enabled the identification of subject-specific motor learning not discernible from conventional kinematic aftereffects. This approach also revealed age-related decline on sensorimotor adaptation in a post-hoc analysis, suggesting that older populations may have limited potential to correct their movements through error-based protocols simply given their age. Further, our detailed EMG characterization provides valuable normative data of muscle activity informing our understanding of the therapeutic effect of split-belt walking.

## 1 Introduction

Humans adapt and learn new movements through interactions with the world, but there have been limited efforts investigating the modulation of muscle activity underlying this motor adaptation. For example, in locomotion, there have been several studies characterizing changes in kinematics (e.g. Reisman et al. 2005), kinetics (Ogawa et al. 2014, Sombric et al. 2018), metabolic cost (Finley et al. 2013, Sánchez et al. 2017), and the perception of movements (Vazquez et al. 2015) that are retained following walking on a split-belts treadmill that moves peoples’ legs at different speeds. However, the studies on muscle activity (Ogawa et al. 2014, Raja et al. 2013, Maclellan et al. 2014) do not provide a detailed characterization of what is adapted in muscle space during and after the novel split-belts situation. It is important to study the adaptation of muscle activity because it provides a distinct and more accurate representation of adjustments in neural commands than conventional biomechanical metrics. Namely, muscle activity is more directly reflective of neural control commands, whereas kinematics and kinetics arise from interactions between those commands and the environment (Collins et al. 2005), and the higher dimension of muscles compared to joints allows distinct muscle coordination to produce the same movements (Bernstein 1967). Therefore, muscle recordings might reveal adjustments of motor commands (Ranganathan and Scheidt 2016) that are obscured by solely characterizing changes in kinematic or kinetic variables.

Muscle activity can be adapted by adjusting planned and corrective motor commands that contribute distinctively to motor adaptation. Planned motor commands are predictive in nature and rely on internal models of the body and the environment (Wolpert et al. 1998). Conversely, corrective motor commands arise during movement execution through a transformation of current sensory information into appropriate actions (Jordan and Rumelhart 1992, Bhushan and Shadmehr 1999). While adaptation of planned movements is slow because it requires repeatedly experiencing a systematic perturbation to update internal models, motor adjustments through corrective actions are rapidly generated immediately after movement disturbances. For example, corrective actions could be strategic adjustments in subsequent movements, such as re-aiming the reaching direction (Morehead et al. 2015, McDougle et al. 2016) or foot landing in walking (Matthis et al. 2017). Corrective commands also include coordinated motor patterns rapidly generated by feedback processes transforming somatosensory input into actions counteracting external disturbances while the movement is ongoing (Crevecoeur and Scott 2013, Chvatal et al. 2011). Some studies presume that rapid changes in motor commands following environmental transitions in walking are merely reactive in nature, reflecting differences in environment-specific walking requirements and distinct from the predictive control processes that are susceptible to sensorimotor recalibration (Reisman et al. 2005, Morton and Bastian 2006, Yokoyama et al. 2018). Conversely, studies on volitional motor control suggest that internal models are available to processes generating both planned and corrective motor commands (Wagner and Smith 2008, Yousif and Diedrichsen 2012). In this study we aim to determine the extent to which corrective responses are indicative of sensorimotor adaptation in locomotion.

We hypothesized that the structure (i.e., motor patterns across multiple muscles) of corrective responses upon removing a novel environment will indicate the motor system’s recalibration. Specifically, that a recalibrated system would adopt the novel situation as a new ‘reference’ such that deviations from it would induce corrective responses of the same structure as those elicited by other novel situations. This is innovative because sensorimotor recalibration is conventionally measured with aftereffects, rather than corrective responses. We defined corrective responses as rapid changes in EMG activity upon transitioning between walking environments or conditions (e.g., legs moving at same speeds vs. different speeds). We tested our hypothesis by analyzing post-adaptation corrective responses, which are those elicited upon transitioning back to the original walking condition following split-belt walking. We considered that these corrective responses would merely reflect changes in the environment if they were numerically opposite to those originally observed when transitioning into the split condition. Conversely, they would be indicative of the motor system’s adapted state if their structure resembled that of corrective responses to transitioning *de novo* to the opposite split environment (i.e., opposite speed difference). This is based on the observation that removal of an altered environment facilitates learning of an opposite environment (Herzfeld et al. 2014b). We further validated our hypothesis by contrasting the structure of corrective responses following short and long durations of the split environment. We expected that the hypothesized structural changes post-adaptation would be exclusively present after the long, but not the short exposure to split-belt walking.

## 2 Materials and Methods

### 2.1 Subjects

A group of 16 healthy subjects of ages ranging between 46 to 78 years old (61+-9.9 y.o., 10 female) participated in the study (see Table 2). We selected an age rage similar to that of stroke survivors (Mozaffarian et al. 2016) because split-belts walking could be exploited to improve the gait of this clinical population (Reisman et al. 2013, Lewek et al. 2017). Therefore, there is an interest in characterizing the possible limitations in the adaptation of motor patterns solely due to the individuals’ age. Their movements and muscle activity were recorded before, during, and after walking on a split-belt treadmill. Subject S01 was excluded from our analyses because this individual failed to follow the instructions during the split-belts walking period, which likely compromised this subject’s motor adaptation. All subjects provided written informed consent prior to participating in the study, which was approved by the Institutional Review Board at our institution, and was in accordance to the declaration of Helsinski.

**Table 1:**
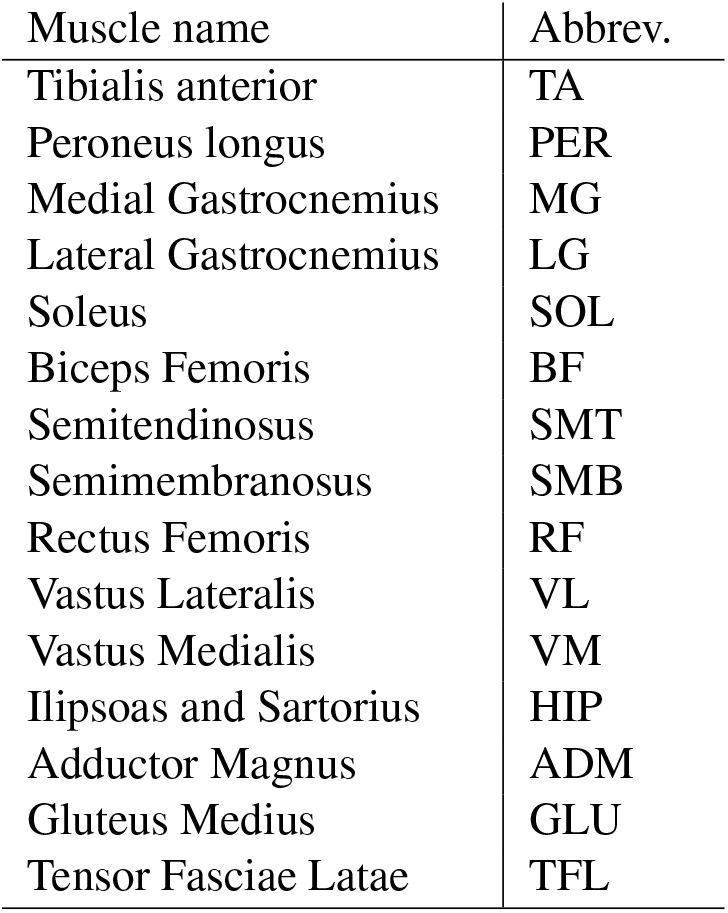
List of recorded muscles & abbreviations.

**Table 2:**
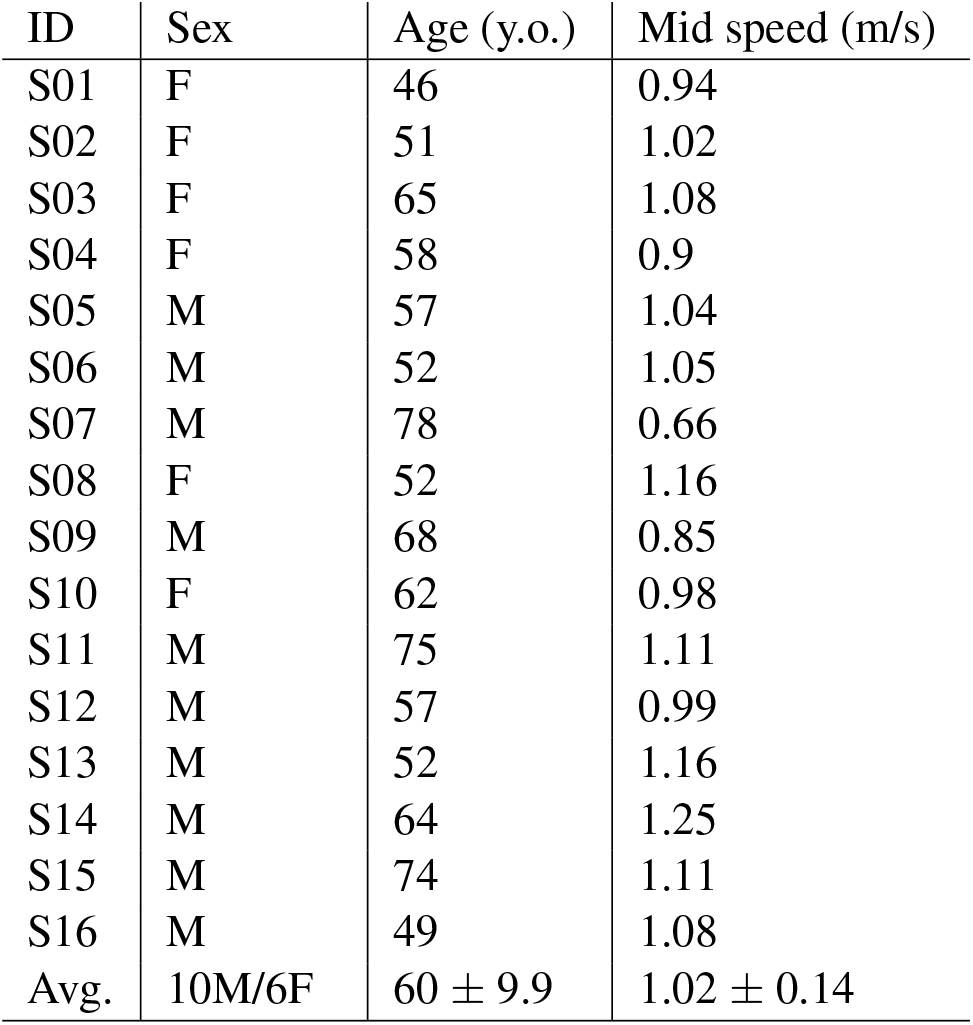
Subject summary. Bottom values indicate mean ± standard deviation.

### 2.2 Experimental Design

We assessed the adaptation and de-adaptation of muscle activity through the split-belts walking adaptation protocol illustrated in Figure 1A. Throughout the experiment, subjects were exposed to three different combinations of belt speeds (walking environments) on a split-belt treadmill (Bertec Corporation, Columbus, OH, USA). Two of these environments were tied-belts walking, in which both belts moved at the same speed, at both a ‘medium’ (self-selected) and a ‘slow’ (33% slower) speed. The third was split-belts walking in which one belt moved at the same slow speed and the other at a ‘fast’ (33% faster than medium) speed, such that the fast belt was moving at twice the speed of the slow one. All subjects experienced the three environments as six different conditions presented in the following order: Slow Walking (50 strides, tied, slow speed), Mid Walking (50 strides, tied, medium speed), Short Exposure (10 strides, split), Baseline (150 strides, tied, medium speed), Long Exposure (900 strides in three blocks of 300, split) and Washout (600 strides in two blocks of 300 or a single 600 block, tied, medium speed). The treadmill was started at the beginning and fully stopped at the end of each block, and speeds did not change while each lock was ongoing. We compared muscle activity throughout the protocol to that of Baseline walking, which was used as a reference. Slow walking was used to quantify speed-dependent modulation of muscle activity during regular treadmill walking. Short and Long Exposure were the only conditions where subjects were exposed to split-belts walking. We selected two exposure durations to dissociate the effect of sensorimotor recalibration from that of sudden changes in walking environment. Namely, the Short Exposure was limited to 10 strides because this is not sufficient to induce sensorimotor recalibration of internal models of walking (e.g. Roemmich and Bastian 2015), whereas the Long Exposure was designed to have 900 strides to ensure muscle activity had reached a steady state. The Long Exposure and Washout conditions were divided into blocks to minimize fatigue and subjects were instructed not to step between blocks to prevent de-adaptation due to unrecorded steps.

**Figure 1:**
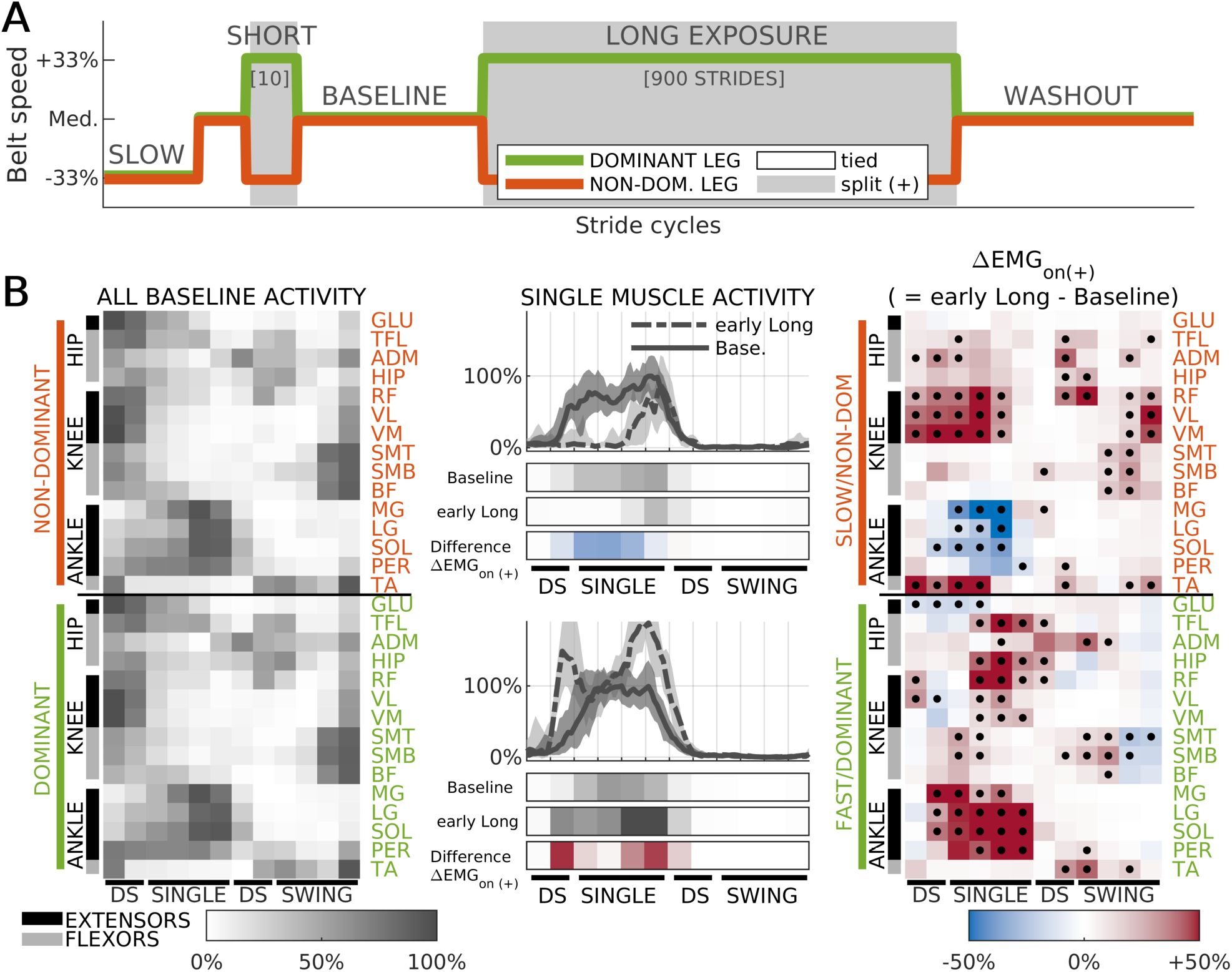
Summary of methods utilized in this study. **(A)** Schedule of belt speeds experienced by all subjects. **(B) Middle column:** sample EMG traces of one muscle (LG) during Baseline (solid) and early Long Exposure (dashed) for a representative subject (S14). Median activity across strides (lines), and the 16-84 percentile range (shaded). Data in traces was processed as described in Methods and further lowpass filtered solely for visualization purposes. Colorbars below the traces represent averaged normalized values during 12 kinematically-aligned phases of the gait cycle (2 for double-support, DS, 4 for single-stance, SINGLE, 2 for second double-support, DS, 4 for swing, SWING, see Methods) for Baseline and early Adaptation (both gray), and the difference between the two (ΔEMG_on(+)_, red indicates increase, blue decrease). The activity of each muscle is aligned to start at ipsilateral heel-strike. Top panels: data for non-dominant/slow leg. Bottom panels: dominant/fast leg. **Left column:** Summary of muscle activity during Baseline walking for all muscles. Median across subjects. Because of the alignment procedure, each column of muscle activity variables is synchronous for all muscles in the non-dominant (top panel) and dominant (bottom panel) legs separately, but not across legs. **Right column:** Summary of change in muscle activity from Baseline to early Adaptation (ΔEMG_on(+)_). Red colors indicate higher levels of activity during early Adaptation, while blue colors indicate lower values. Median across subjects. Black dots indicate significant differences from 0. P-value threshold *p* = 0.035

The ‘medium’ walking speeds were determined in a self-selected way to ensure subjects from all ages could complete the entire protocol. The self-selected speed was obtained by first averaging each subject’s speed when walking over ground in a 50-meter hallway during a 6-min walking test (Rikli and Jones 1998) and then subtracting 0.35*m/s*, which resulted in a comfortable walking speed on a treadmill based on pilot data in older adults (> 65 yrs). This resulted in ‘medium’ walking speeds of 0.72 ± 0.26*m/s* (mean ± standard deviation) across the population. Medium walking speeds for each individual can be found in Table 2. The ‘slow’ speed was defined as 66.6% of the ‘medium’ speed, and the ‘fast’ speed as 133.3% of the same. In this way, the average belt-speed during split-belts walking matched that of Baseline and Washout, and the belt-speed ratio during split-belts walking was 2:1. When walking in the split-belts environment, the dominant leg (self-reported leg used to kick a ball) was always walking faster. Thus, we refer to the dominant leg as the fast leg and the non-dominant one as the slow leg throughout the text.

Safety measures were designed such that participants from older ages could complete the study. First, all subjects wore a harness that only provided weight support in the event of falling but not during walking or standing. Also, subjects were told a few seconds in advance that they could hold on to a handrail (directly located in front of them) whenever a condition or block started or finished. Subjects were encouraged to let go of the handrail as soon as they felt comfortable doing so to minimize the effect of external support on muscle recordings. Finally, we monitored subjects’ heart-rate continuously and blood-pressure during the rest breaks to prevent over exertion in any of the participants.

### 2.3 Data analysis

#### Acquisition & pre-processing

We collected electromyographic (EMG) signals, kinematics, and kinetic data to characterize subjects’ behavior. Surface EMG signals from 15 muscles on each leg were recorded for all subjects (see Table 1 for full list and abbreviations) at 2000Hz using a Delsys Trigno System (Delsys Inc., Natick, MA, USA). Signals were highpass-filtered to remove undesired movement artifacts and then rectified. We used a 2nd order Butterworth filter (dual-pass) with a cutoff frequency of 30Hz, which resulted in 80 dB/dec attenuation and zero-lag (Merletti and Parker 2004). Unlike other studies (e.g. Torres-Oviedo and Ting 2007), we did not apply a subsequent lowpass filter following rectification as we did not require the EMG envelope for our analysis (see below). Kinematic data was collected at 100 Hz with a passive motion analysis system (Vicon Motion Systems, Oxford, UK). Movements were recorded by placing reflective markers bilaterally on bony landmarks at the ankle (i.e., lateral malleolus) and the hip (i.e., greater trochanter). Ground reaction forces were recorded with an instrumented split-belt treadmill (Bertec Corporation, Columbus, OH, USA) and sampled at 1000Hz. Forces along the axis of gravity (*F_z_*) were used to determine when the foot landed (i.e., heel-strike: *F_z_* > 10N) or was lifted off the ground (i.e., toe-off: *F_z_* < 10*N*).

#### EMG parameters

EMG activity was time-aligned, binned, and amplitude-normalized to allow for comparisons across epochs, muscles, and subjects (Figure 1B, middle column). First, filtered EMG activity was divided in intervals of the gait cycle aligned to gait events to focus on changes in muscle activity within the gait cycle, rather than on changes due to differences in timing of the gait cycle across the distinct walking conditions (Dietz et al. 1994, Reisman et al. 2005). More specifically, we divided the gait cycle of each leg into 4 intervals according to well defined gait phases (Perry and Burnfield 2010): first double-support (DS, from ipsilateral heel-strike to contralateral toe-off), single-stance (SINGLE, from contralateral toe-off to contralateral heel-strike), second double support (DS, from contralateral heel-strike to ipsilateral toe-off), and swing (SWING, from ipsilateral toe-off to ipsilateral heel-strike). In order to achieve better temporal resolution, each of these 4 intervals were further divided. The double-support phases were divided into two equal sub-intervals, and the swing and single-stance phases divided into four equal sub-intervals, yielding 12 intervals for each gait cycle. The average duration of these sub-intervals ranged between 75 and 120 ms. Specific timing for each interval throughout the different epochs of the study are presented in Table 3. Muscle activity amplitude was averaged in time for each of these sub-intervals for every stride and muscle resulting in 360 muscle activity variables per subject per stride cycle: 12 sub-intervals × 30 muscles. These variables were then amplitude-normalized to be able to compare measurements across subjects and muscles. For each subject and muscle, we first computed the mean activity for each sub-interval across the last 40 strides (i.e., steady state) of Baseline walking. Then each muscle’s activity was scaled and shifted in magnitude such that the least and most active sub-intervals for that subject and muscle during those last 40 strides of Baseline took the values of 0 and 100%, respectively. The same scaling and shifting was used for activity during all conditions of our experiment so that muscle activity is represented as a percentage of the maximum Baseline activity. Because of this, all muscle activity variables used in this study are dimensionless. This scaling allowed us to aggregate subjects and compare effect sizes across muscles even when recorded EMG amplitudes were very different because of sensor placement or underlying tissue properties. We present in Figure 1B sample traces of time-aligned and amplitude-normalized EMG activity for one sample muscle averaged across gait cycles for late Baseline and early Long Exposure (Figure 1B, middle column). The gray rows indicate the results of the time-binning for two experimental epochs: late Baseline and early Long Exposure. The rows colored in red and blue represent the difference between these two epochs. The left column of Figure 1B shows the resulting time-binned and amplitude-normalized activity for all muscles averaged across subjects and strides during late Baseline. The right column of Figure 1B shows the difference in time-binned and amplitude-normalized activity between early Long Exposure and late Baseline. These changes in muscle activity upon the introduction of the split-belts perturbation (ΔEMG_on(+)_) was used to characterize the structure of corrective responses following the Short Exposure and Long Exposure of the split-belts condition (as explained in detail in section 2.4). This coordinated pattern of corrective responses (ΔEMG_on(+)_) to the novel walking environment was very similar to balance-like responses previously reported (Torres-Oviedo and Ting 2007, Chvatal and Ting 2012, Safavynia and Ting 2012, de Kam et al. 2017). For example, the ΔEMG_on(+)_ activity of the low leg during SINGLE initially showed increased activity of anterior muscles (TA, quads -RF, VM, and VL-, and hip flexors -HIP, ADM, TFL-) and reduced the activity of calf muscles (MG, LG, SOL), as has been observed in balance responses when the body falls backward (Tang and Woollacott 1999, Chvatal et al. 2011, Chvatal and Ting 2012).

**Table 3:** Mean time elapsed between relevant kinematic events (heel-strikes, HS, and toe-offs, TO) during the different epochs of the experiment. Intervals are presented in order of occurrence in the gait cycle, starting at the fast leg’s heel-strike (FHS). Median ± interquartile range values across participants.

| Epoch | DS1 [FHS-STO] (ms) | swing SLOW (ms) | DS2 [SHS-FTO] (ms) | swing FAST (ms) |
| --- | --- | --- | --- | --- |
| Baseline | 168 ± 12 | 393 ± 59 | 166 ± 29 | 383 ± 43 |
| early Long | 177 ± 46 | 314 ± 64 | 135 ± 34 | 458 ± 128 |
| late Long | 176 ± 21 | 329 ± 38 | 161 ± 30 | 473 ± 114 |
| early Wash. | 150 ± 35 | 358 ± 91 | 188 ± 32 | 372 ± 69 |
| late Wash. | 187 ± 24 | 405 ± 42 | 181 ± 33 | 394 ± 52 |

#### Kinematic parameters

The adaptation of movements was characterized with step-length asymmetry, which is a metric known to change during and after split-belts walking (Reisman et al. 2005). We computed step-length asymmetry on each stride cycle by calculating the difference in step lengths (i.e., ankle to ankle distance at foot landing) for two consecutive steps taken with the fast and slow leg. This difference was normalized by the sum of step lengths to obtain a measure that was a proportion of each subjects’ step sizes. A zero step-length asymmetry value indicated that steps lengths were even, negative values indicated that the (non-dominant) leg walking on the slow belt was taking longer steps than the (dominant) one on the fast belt and viceversa for positive values. We also computed body displacement with respect to the foot in contact with the ground during the stance phase for each leg. This was done to interpret the changes in muscle activity upon transitions between tied and split conditions. Body displacement was computed as the anterior-posterior distance between the middle of the hip markers (greater trochanter) and the ankle from ipsilateral heel-strike to contra-lateral heel-strike. Kinematic data was time-aligned to kinematic events (heel-strikes and toe-offs) in the same way as EMG data.

#### Epochs of interest

For our analysis, we considered behavioral measurements only at the beginning (‘early’) and end (‘late’) of each experimental condition (epochs). ‘Early’ epochs were characterized by the average of 5 strides and ‘late’ epochs by the average of 40 strides. Recordings during the very first and last stride of each condition were excluded to eliminate effects linked to the treadmill’s starting and stopping. For example, early Long Exposure consisted of the mean activity for strides 2-6, such that 5 strides were considered but the first one was excluded. For Short Exposure we used all available strides after discarding the first and last (a total of 8) instead of identifying separate early and late epochs.

### 2.4 Characterizing corrective responses to sudden changes in the environment

We proposed to study the structure of corrective responses to sudden changes in walking conditions (Figure 2A). Corrective responses are defined to be the rapid changes in motor output immediately after a transition in the walking environment (i.e., within 5 steps after the speed transition is experienced). Per our definition, corrective motor responses include modulation of muscle activity that occurs at different latencies after a sudden change in the environment, which may or may not be voluntary (i.e., short- and long-latency reflexes, subsequent voluntary responses, and changes in strategy) (Horak et al. 1990). We quantified corrective responses as the difference in EMG activity following changes in walking conditions, that is averaged EMG activity in the epoch after the transition (i.e., *EMG*_after_) with respect to the epoch before it (i.e., *EMG*_before_, see description of epochs above). For example, in the Baseline to Long Exposure transition *EMG*_after_ represents the average of the first 5 strides of Long Exposure and *EMG*_before_ the average of the last 40 strides of Baseline. Thus:

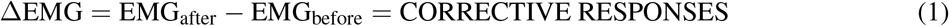

**Figure 2:**
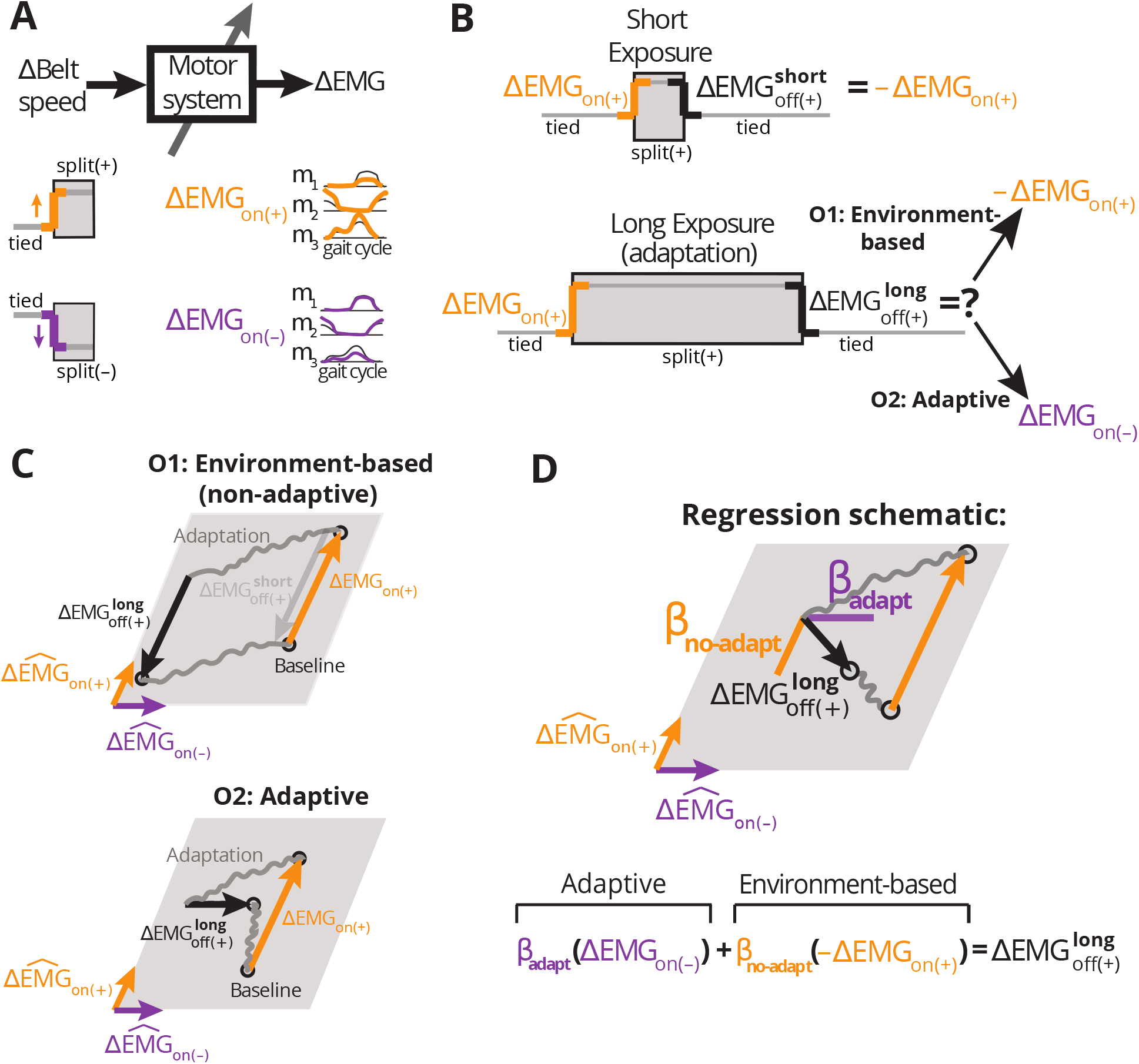
Schematic of the expected corrective responses following short and long exposures to split-belts walking. **(A)** Schematic of the input-output relation (system) of study. We consider belt-speeds (the walking environment) as the input to the system and motor commands (as measured by EMG activity) as its output. We specifically proved rapid EMG changes in response to sudden transitions in the walking environment. Hypothetical EMG patterns in response to an ‘on (+)’ and ‘on (–)’ transition are illustrated in yellow and purple, respectively. In the ‘on (+)’ transition the dominant leg walks unexpectedly faster than the non-dominant one and vice versa for the ‘on (–)’ transition. **(B)** We contrasted corrective responses upon removal of the (+) environment following either a short 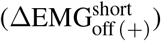 or a long 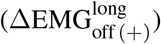 exposure duration. In the case of a short exposure (top) we expect muscle activity to return to the activation pattern before the introduction of the split-belts environment. In the case of a long exposure (bottom) corrective responses could either remain the same as those following a short exposure (i.e., non-adaptive, O1); or be adaptive, exhibiting a structure similar to that in the ‘off (–)’ transition (O2). **(C)** Schematics of hypothetical patterns of activity in muscle space under the environment-based (O1) and adaptive (O2) alternative outcomes. We present a two-dimensional muscle space for illustration purposes, but we characterized muscle patterns in a 360-dimensional muscle space. A point in this space represents a pattern of activity across all muscles, whereas colored arrows represent changes in muscle activity from one activation pattern to another upon an environmental transition. EMG changes over the course of adaptation and washout periods (gray) were not investigated. Under O1, 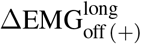 (black) and ΔEMG_on(+)_ (yellow) are expected to be numerically opposite. Under O2, 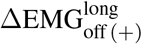 is expected to be equal to ΔEMG_on(–)_ (purple). **(D)** Schematic and equation of the regression model used to quantify the structure of 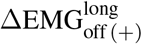. *β*_adapt_ quantifies the similarity to responses expected under O2, whereas *β*_no-adapt_ to those under O1.

We used the following subscripts to define the corrective responses (ΔEMG) to different environmental transitions: ‘on’ and ‘off’ indicate ΔEMG when the split environment is introduced or removed, respectively. ‘(+)’ and ‘(–)’ indicate the particular split environment that was introduced or removed. Specifically, ‘(+)’ arbitrarily indicates that the belt under the dominant leg was moving faster than the one under the non-dominant leg (i.e. the environment used in this experiment) and ‘(–)’ indicated the opposite situation (i.e., the non-dominant leg moving faster than the dominant one, not tested here). While all four combinations of ‘on’ or ‘off’ and ‘(+)’ or ‘(–)’ may happen, only three of them are relevant to our study: the introduction of the two split-belt environments ‘on (+)’ and ‘on (–)’, and the removal of the ‘(+)’ environment: ‘off (+)’. Lastly, we used ‘short’ and ‘long’ superscripts to distinguish between exposure durations to the novel split environment (i.e., 10 strides vs. 900 strides respectively), for ‘off (+)’ transitions only.

In order to test our hypothesis, we analyzed the structure of corrective responses upon the removal of a split-belts walking environment (ΔEMG_of (+)_, Figure 2A). Specifically, we hypothesized that the structure of those responses may change because of sensorimotor recalibration (Figure 2B). We analyzed the structure through the following regression model illustrated in Figure 2D:

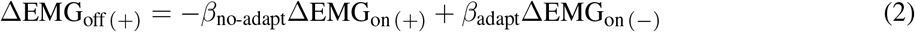

Where ΔEMG_on(+)_ and ΔEMG_on(–)_ represent the corrective responses observed when the ‘(+)’ or the ‘(–)’ environment are initially introduced, respectively (a cartoon of these responses is displayed in Figure 2B).

Upon the Long Exposure (900 strides), when we expect subjects to experience sensorimotor recalibration, we considered two potential outcomes (Figure 2C): solely environment-dependent (non-adaptive) corrective responses (O1), and adaptive corrective responses (O2). Under the first possibility, we expect corrective responses to the ‘off’ transition would be numerically opposite to those upon introduction (‘on’) of the novel environment, as expected when switching back and forth between two environment-specific motor patterns (A to B vs. B to A). Formally expressing O1:

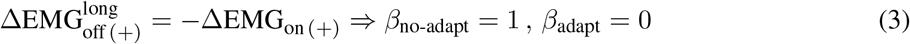

Under the second alternative (O2) the motor system learns that split-belts walking is the “new normal”. Consequently, removal of the ‘(+)’ environment would be processed as a perturbation opposite to the one originally experienced (i.e. it would be equivalent to a ‘on (–)’ transition). This would be consistent with previous work reporting that the removal of altered environmental dynamics is in itself a perturbation (Herzfeld et al. 2014b) and that corrective responses are adapted through experience to an altered environment even in the absence of feedback-specific learning opportunities (Wagner and Smith 2008, Yousif and Diedrichsen 2012). Formally expressing O2:

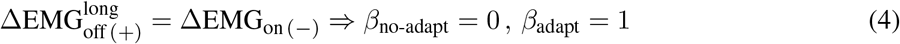

In sum, our general regression model quantified concurrently the extent to which the structure of corrective responses post-adaptation were environment-dependent (O1) and adaptive (O2) with *β*_no-adapt_ and *β*_adapt_, respectively (Figure 2D).

Note that *β* coefficients might be smaller than expected because of differences in the predicted structure or because magnitude differences between ΔEMG_on(+)_ and ΔEMG_off (+)_. To dissociate the effect of magnitude from that of structure of corrective responses, we performed a secondary analysis in which we tested the alignment of either the ΔEMG_off (+)_ vector and the vectors –ΔEMG_on(+)_ and ΔEMG_on (–)_ by computing the cosine of the angles they form in the 360-dimensional space. This analysis is similar to the regression analysis, but is insensitive to differences in the magnitude of corrective responses. Under (O1) we expect 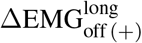 to be aligned (cosine of 1) to –ΔEMG_on(+)_, and under (O2) we expect it to be aligned to ΔEMG_on (–)_. All analyses were performed for both individual and group averaged patterns of activity.

As validation, we performed the same analyses following the Short Exposure (10 strides) to the ‘(+)’ environment, when we expect subjects will not undergo sensorimotor recalibration. In this case, we expect corrective responses to be strictly consistent with the first alternative (O1, Figure 2C, top).

#### Inferring corrective responses to an opposite transition

We did not directly measure ΔEMG_on (–)_ to avoid exposing subjects to multiple environmental transitions. Instead, we infer these responses by exploiting the symmetry of the transition between the two legs. Assuming that leg dominance plays no role, the only difference between the ‘(+)’ and ‘(–)’ environments is which leg moves faster than the other. In other words, the environments are mirror images of one another. Thus, we expect the corrective responses ΔEMG_on (–)_ to be the same as ΔEMG_on(+)_ but transposing the activity for the two legs. That is, upon a ‘(–)’ transition we expect to see in the dominant leg the same responses that are observed for the non-dominant leg in the ‘(+)’ transition and vice versa. It is worth pointing out that our regression analyses assumes that ΔEMG_on (–)_ and – ΔEMG_on(+)_ muscle vectors are different from each other in muscle space (i.e., not colinear). Because of how we estimate ΔEMG_on (–)_, this implies that initial muscle responses (ΔEMG_on(+)_) should not be anti-symmetric (i.e., same group of muscles increasing in a leg, decrease in the other one). This assumption was confirmed empirically, as the cosine of the angle formed by the two vectors was –0.13 ± 0.38 (median ± inter-quartile range across subjects).

### 2.5 Statistical analysis

#### Differences in muscle activity across epochs

Significant changes in muscle activity variables between any pair of epochs were determined by using the Wilcoxon signed-rank test (i.e., non-parametric analogue of a paired t-test) on each of the 360 muscle activity parameters. Effect sizes for the population were computed using median values across subjects since this is a measure less susceptible to outliers. All tests were twotailed and the null hypothesis was that there was no difference in the activity between the two epochs being compared. In order to limit false discoveries given the large number of comparisons performed, we used the Benjamini-Krieger-Yekuteli two-stage procedure to control the False Discovery Rate (FDR) (Benjamini and Hochberg 1995, Benjamini et al. 2006, Kass et al. 2016). We set the acceptable FDR to be 0.05 (i.e. that no more than 1 out of 20 significant findings will be false positives in expectation). Threshold (critical) p-values from the FDR procedure are reported for each application. Solely for visualization purposes, we choose to indicate only the results that are significant after the FDR-controlling procedure and that correspond to an absolute effect size larger than 10% (i.e., 10% of the maximum Baseline activity for that muscle, see the procedure for amplitude-normalization of muscle activity above). This was done to avoid showing statistically significant but small, and presumably meaningless, differences.

#### Structure of corrective responses

The linear regressions characterizing the structure of feedback-generated activity were performed using Matlab’s *fitlm* function and computing (Pearson’s) *R*^2^ values that were uncentered, given that our regression model did not include intercept terms. We compared the regressors obtained for data following the Short and Long Exposure conditions ΔEMG_of (+)_ using a two-tailed paired t-test. We report p-values, as well as mean changes and well as Cohen’s d for effect size.

#### Analysis of inter-subject variability

We conducted post-hoc regression analyses to determine if either age or walking speed could explain the large inter-subject variability that we observed in the regression coefficients. We focused on these subject-specific features because they exhibited large ranges in our cohort that could have impacted our results. We also studied the association of age with the magnitude of muscle activity aftereffects, corrective responses and step-length asymmetry. Magnitude of muscle activity variables (aftereffects, corrective responses) was computed as the euclidean norm of the relevant 360-dimensional vectors (e.g., ║*EMG*_early Wash._ – *EMG*_Baseline_║ for aftereffects, ║ΔEMG_on(+)_║ for ‘on’ corrective responses). For all these analyses we used Spearman’s correlations, which is a non-parametric alternative to Pearson’s correlation, because it is more robust to potential outliers (Rousselet and Pernet 2012). The correlation value (*ρ*) and the corresponding p-value were presented. For the relation between regressors and age, Pearson’s correlations are also presented as a reference. All results from the inter-subject analysis are reported in section 3.3 of the results.

### 2.6 Data and code availability

All data and code used for this study are available at doi:10.6084/m9.figshare.c.4423772

## 3 Results

### 3.1 Modulation of muscle activity during late Long Exposure is not a direct reflection of each leg’s walking speed

Muscle activity during late Long Exposure was not simply regulated as a function of the speed at which each leg moved in the split condition. Overall we observed an anti-symmetric (i.e., similar magnitude but opposite sign) pattern of modulation of muscle activity of the two legs during late Long Exposure with respect to Baseline (Figure 3B). That is, we found that if a group of muscles increased activity on one side, the same group decreased activity on the other one. Interestingly, this opposing modulation across legs was not merely determined by the belt’s speed under each leg (i.e. ipsilateral walking speed), with all muscles on one leg increasing activity and all muscles on the other leg decreasing it. Should this have been the case, one would expect reduced activity of the slow leg with respect to Baseline (medium speed) as seen in slow walking (Figure 3A), and increased on the fast leg (Den Otter et al. 2004, Dietz et al. 1994). However, this type of modulation was only observed in distal muscles (MG, LG, SOL, PER, TA) and in the fast leg’s hip flexors (HIP, TFL, ADM), but not in most of the proximal muscles (SMB, SMT, BF, RF, VM, VL) and (ADM, TFL) of the slow leg, which increased in the leg walking slow and decreased in the leg walking fast. We found little similarities in the modulation of muscle activity between late Long Exposure and early Washout. We indicate with black outlines in Figure 3B and 3C the muscles that do show the same modulation sign in both. This suggests that the muscle activity that gives rise to behavioral aftereffects cannot simply be inferred from the steady-state motor patterns observed in an altered environment. In sum, muscle activity for split-belts walking highlights the interlimb nature of locomotion because it exhibited an interaction between muscle group (i.e., distal vs. proximal) and walking speed, rather than individual leg modulation based on speed.

**Figure 3:**
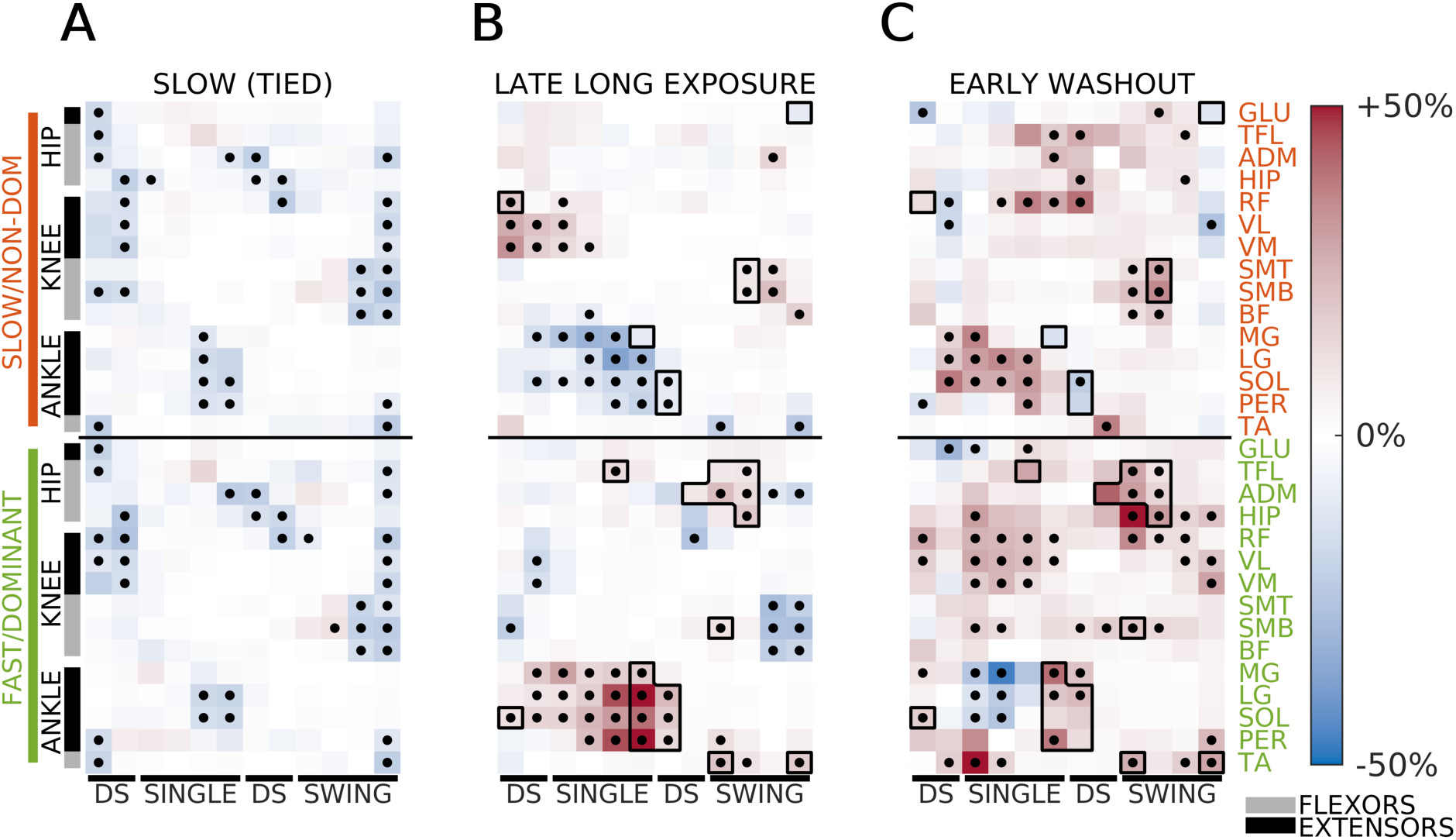
Steady-state muscle activity during slow walking, late split-belts walking, and aftereffects. Panels reflect differences in muscle activity the three epochs with respect to the reference (Baseline) condition. Colormap reflects effect size and dots indicate FDR controlled significant differences (see Methods). Muscle activation variables were displayed starting with the ipsilateral heel-strike. **(A)** Muscle activity modulation during Slow (tied-belts) walking. Most muscle-phases show reduction of activity, consistent with a monotonic link between walking speed and muscle activity amplitude. P-value threshold: *p* = 0.022. **(B)** Muscle activity modulation during late Adaptation. Broadly, patterns of activity are anti-symmetric, with groups of muscles increasing activity in one leg and decreasing contralaterally. P-value threshold: *p* = 0.022. Differences between the slow leg’s activity at late Long Exposure (panel B, top) and Slow (tied) baseline (panel A, top) illustrate that split-belts patterns do not match the expectation from simple ipsilateral speed modulation. **(C)** Muscle activity modulation during early Washout with respect to Baseline. Black dots indicate significance. P-value threshold: *p* = 0.026. Few similarities are found between the steady-state activity during late Long Exposure and observed aftereffects. Black outlines indicate the muscle-intervals with activity changes of at least 10% and the same sign (i.e. increase or decrease with respect to Baseline) for both late Long Exposure and early Washout.

### 3.2 Sensorimotor recalibration after prolonged exposure to split-belts walking is indicated by the structure of corrective responses

To quantitatively characterize the structure of corrective responses upon removal of the novel split-belts environment, we used a regression model (ΔEMG_off(+)_ = –*β*_no-adapt_ΔEMG_on(+)_ + *β*_adapt_ΔEMG_on(–)_) (Figure 4). If corrective responses were purely environment-dependent (O1) we expect that muscle patterns upon introduction and removal of the split condition would be numerically opposite (i.e., ΔEMG_off(+)_ = –ΔEMG_on(+)_, thus *β*_adapt_ = 0, *β*_no-adapt_ = 1, Figure 4A), whereas under the adaptive possibility (O2), it would be equivalent to experiencing the introduction of the opposite perturbation (i.e., ΔEMG_off (+)_ = ΔEMG_on (–)_, thus *β*_adapt_ = 1, *β*_no-adapt_ = 0, Figure 4B).

**Figure 4:**
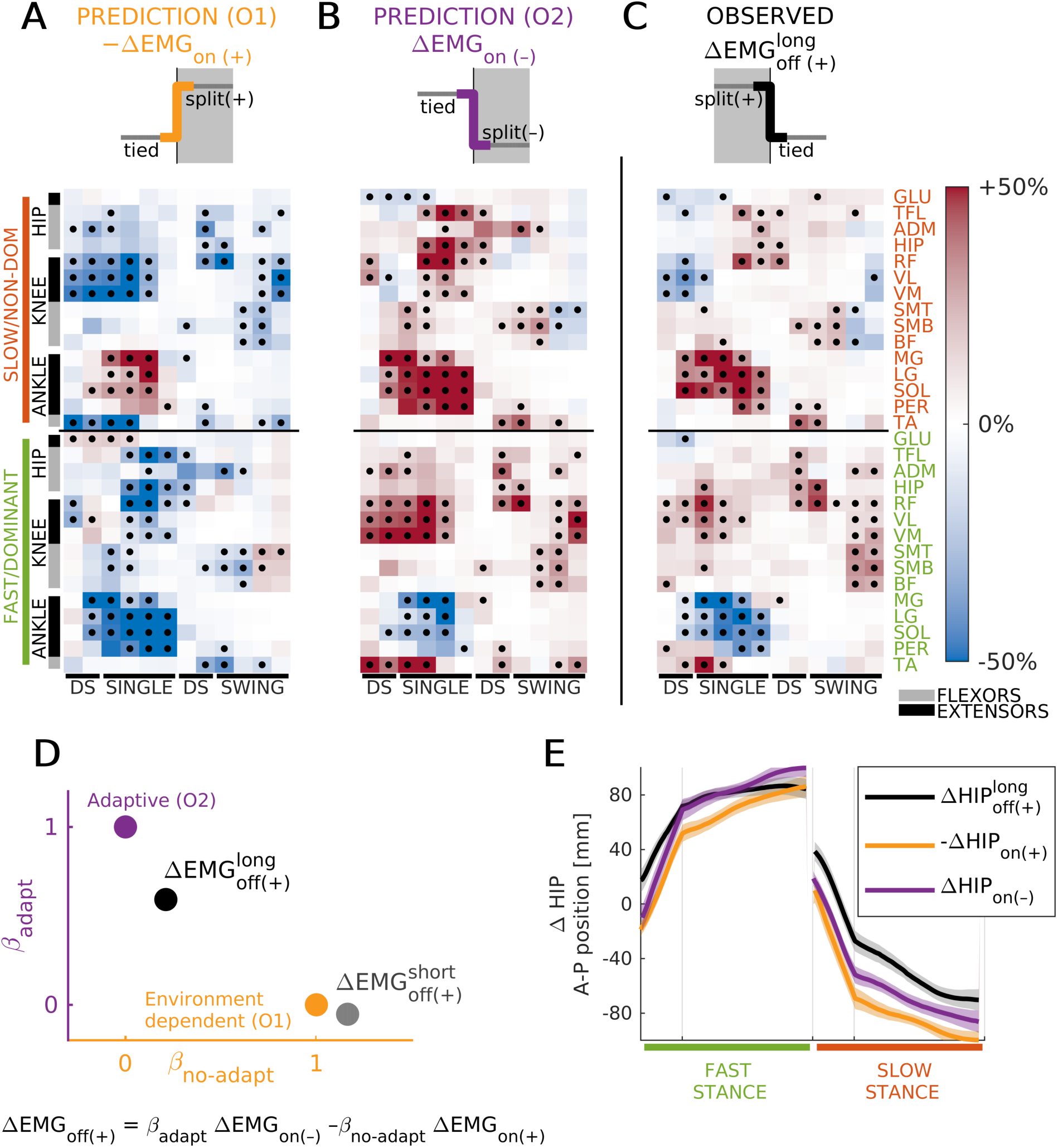
Corrective responses were adapted following a long exposure to a split-belts walking environment. **(A,B)** Expected corrective responses elicited by the ‘off’ transition under the environment-based (O1, panel A) and adaptive (O2, panel B) cases. Data (in color) and significance (black dots) were derived from the observed corrective responses upon the introduction of the ‘(+)’ walking environment (Figure 1B, right column), by either taking the numerical opposite (O1) or by transposing leg activity (O2). For more details, see 2. **(C)** Actual corrective muscle activity responses upon removal of the ‘(+)’ environment (i.e., ‘off’ transition). Black dots indicate significant changes in activity following FDR correction. P-value threshold: *p* = 0.035. **(D)** Quantification of corrective responses’ structure upon removal of the ‘(+)’ environment following long (black) and short (gray) exposure durations. As expected, responses following the short exposure displayed environment-based structure. In contrast, those following a long exposure appeared as if removal of the novel ‘(+)’ environment was equivalent to introducing a novel ‘(–)’ environment. **(E)** Changes in anterior-posterior hip position (with respect to stance foot) following the long exposure (black). Expected changes under O1 (yellow) and O2 (purple) are also illustrated. These were computed in the same way as EMG factors displayed in A and B. The similarity across all traces indicates that it is impossible to characterize the adaptive and environment-based (non-adaptive) nature of corrective responses solely from a global measure of body position.

For the transition following the Long Exposure period, our regression analyses were consistent with adaptive corrective responses (O2) when using both group averaged activity (Figure 4D, black dot: CI for *β*_adapt_ = [.543, .641], *p* = 5.1 × 10^−76^, *β*_no-adapt_ = [.163, .26], *p* = 4.2 × 10^−16^, *R*^2^ = 0.662), and muscle activity for each individual (Figure 5A, small magenta dots; median ± inter-quartile range for *β*_adapt_ = 0.437 ± 0.112, *β*_no-adapt_ = 0.15 ± 0.317, and *R*^2^ = 0.297 ± 0.167). Importantly, our results were not dependent on the number of strides used to quantify corrective responses. For example, similar results were obtained when using 15 strides, rather than 5, but with slightly larger regression coefficients and *R*^2^ (e.g., group averaged activity with 15 strides: CI for *β*_adapt_ = [.568, .677], *β*_no-adapt_ = [.234, .343], *R*^2^ = 0.642). The same qualitative, although noisier, results are also obtained if we use a single stride for the analysis.

**Figure 5:**
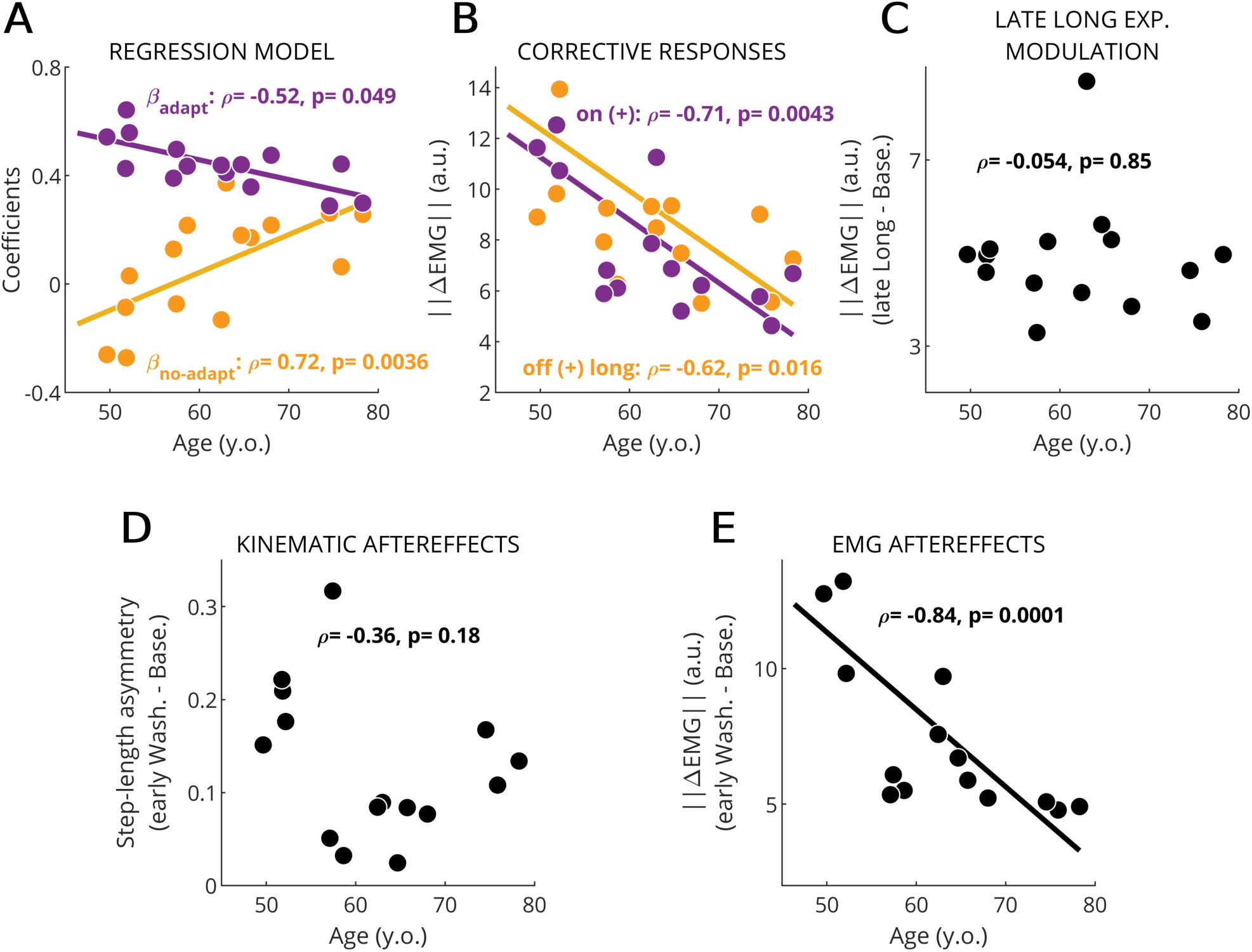
Age modulated EMG-based learning measures, but not kinematic ones. Single dots represent values for one subject. Spearman’s correlation coefficients (*ρ*) and p-values (*p*) are presented on the legend. The best line fit of the dependent variable onto age is only displayed when significant (*p* ≤ 0.05). **(A)** Regression coefficients from model quantifying the structure of EMG corrective responses. (purple: *β*_adapt_, yellow: *β*_no-adapt_). Both regression coefficients were significantly correlated with age. **(B)** Magnitude of corrective activity following the introduction (║ΔEMG_on(+)_║) and removal 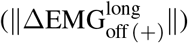 of the ‘(+)’ environment. We observed a significant effect of age at both transitions. **(C)** Magnitude of steady-state changes in muscle activity during late Long Exposure. No correlation to age was found. This confirms that older subjects were able to modulate muscle activity as much as healthy subjects. **(D)** Step-length asymmetry aftereffects were also not correlated with age. **(E)** Size of muscle activity modulation during early Washout (aftereffects). Aftereffects were correlated with age. This suggests EMG-based measures of learning were more sensitive than kinematic-based ones.

The adaptive nature of corrective responses was further supported with a secondary analysis that compared the structural similarities between 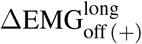 and the regression factors ΔEMG_on(–)_ and – ΔEMG_on(+)_. Specifically, we computed the cosine of the angle between 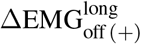 and each of the regression factors. We did this complementary analysis because it was not affected by the magnitude differences between corrective response to the ‘on’ and ‘off’ transitions (i.e., the magnitude of 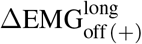 was 81.6% of the ΔEMG_on(+)_ magnitude for individual subjects and 74.8% for group median data). We found that 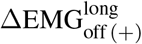 and ΔEMG_on(–)_ were more aligned when using group median activity (cosine of 0.778) and individual data (median ± inter-quartile range: 0.521 ± 0.175) than 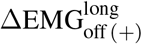 and –ΔEMG_on(+)_ for group (cosine of 0.361) and individual data (median ± inter-quartile range: 0.19 ± 0.595).

Our regression and cosine-based results are qualitatively supported by the remarkable similarity between the 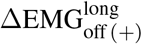 and ΔEMG_on(–)_ (Fig. 4B vs. Fig. 4C), which was predicted under O2 (Fig. 4B, top half). In contrast, the adaptive nature of corrective responses could not be observed in the analysis of body motion (Figure 4E). Specifically, the changes in body position during each leg’s stance phase (when the leg is in contact with the ground) were the same upon removal of the (+) environment (ΔHIP_off (+)_) and the body motion from the two alternatives considered (ΔHIP_on(–)_ and -ΔHIP_on(+)_).

Lastly, to confirm that these patterns of muscle activity arise from a learning-dependent process, and not simply due to removal of the split environment, we ran the same regressions on the split-to-tied transition following the Short Exposure condition (i.e., we used the exact same regression factors as in the analysis for the Long Exposure period). In this condition subjects did not have time to adapt, so we expected to observe changes in EMG activity consistent with environment-dependent transitions (O1). Indeed, that is what we found with our regression model applied to the transition after the Short Exposure 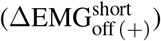. This was true for both the group-averaged activity (Figure 4D, gray dot; CI for *β*_adapt_ = [–.106, –.0002], *p* = 0.049, *β*_no-adapt_ = [1.11,1.22], *p* = 5.9 × 10^−144^, *R*^2^ = 0.840) and the activity of individual subjects (data not shown) (median ± inter-quartile range for *β*_adapt_ = 0.122 ± 0.316, for *β*_no-adapt_ = 0.728 ± 0.471, and for *R*^2^ = 0.465 ± 0.224). In addition, our cosine-based analysis returned values of 0.170 and 0.039 ± 0.336 for the group and individual vectors to ΔEMG_on (–)_ respectively, and 0.919 and 0.671 ± 0.166 for vectors to –ΔEMG_on(+)_. In sum, we found a strong dissociation of the structure of corrective responses and exposure duration: following the Short Exposure condition, corrective muscle activity can be modeled as environment-dependent, whereas following the Long Exposure the corrective responses are better explained by considering that they are adapted. Namely, comparison of the individual coefficients indicated that every subject has a higher *β*_adapt_ and smaller *β*_no-adapt_ after the Long than the Short Exposure, with a median change of 0.40 and –0.643 and Cohen’s d of 1.65 and –2.93, respectively (Long vs. Short: *β*_adapt_: *p* = 1.87 × 10^−6^ and *β*_no-adapt_: *p* = 3.08 × 10^−11^). Consistently, every subject’s corrective responses were more aligned to ΔEMG_on (–)_ and less to –ΔEMG_on(+)_ following the Long Exposure compared to the Short one, with a mean cosine change of 0.558 and –0.539, and corresponding Cohen’s *d* of 1.84 and –2.72 respectively (Long vs. Short paired t-test: *p* = 5.15 × 10^−6^ and *p* = 4.89 × 10^−8^). Taken together, we found that the structure of corrective muscle activity reflected the recalibration of the motor system during adaptation.

### 3.3 Corrective responses revealed age-related decline of sensorimotor adaptation

Regression analysis of inter-subject variability revealed that older adults exhibited less adaptation of corrective responses. Namely, *β*_adapt_ and *β*_no-adapt_ were associated to subjects’ age (*β*_adapt_ vs age: Spearman’s *ρ* = –0.52, *p* = 0.049, Pearson’s *R*^2^ = 0.511, *p* = 0.00275 and *β*_no-adapt_ vs. age: *ρ* = 0.72, *p* = 0.0036, *R*^2^ = 0.423, *p* = 0.0086, Figure 5A) with older subjects showing smaller *β*_adapt_ and larger *β*_no-adapt_. This indicates that corrective responses in older adults were less adapted (and more environment-driven) compared to younger adults. We also observed that the magnitude of corrective responses was smaller in older adults (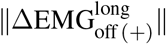: *ρ* = –0.71, *p* = 0.0043; ║ΔEMG_on(+)_║: *ρ* = –0.62, *p* = 0.016; Figure 5B). These smaller responses in older individuals could make it more difficult to identify the structure of corrective responses because of the reduced signal to noise ratio, possibly leading to biased or noisy estimates of *β*_adapt_ and *β*_no-adapt_. To discard this possibility, we correlated the *R*^2^ of the fitted models for each individual with age. We found no effects of age (*ρ* = –0.51, *R*^2^ = 0.25, *p* = 0.058, not shown), meaning that the regression model applied to individual data captured comparable levels of variance regardless of subjects’ age. Taken together, we observed an age-mediated difference in the extent of adaptation of corrective muscle activity, suggesting an age-mediated decline of learning processes updating corrective responses. This limited learning capacity in older individuals was further supported by the negative association between age and the magnitude of EMG aftereffects (║EMG_early Wash._ – EMG_Base._║, *ρ* = –0.84, *p* = 0.0001, Figure 5E): the older the subjects, the smaller the aftereffects. This happened despite the similar magnitude of muscle activity during late Long Exposure across individuals (║EMG_lateLong E._ – EMG_Base._║ vs. age: *ρ* = –0.054, *p* = 0.85, Figure 5C). Importantly, walking speed, which naturally alters muscle activity, was not associated to the magnitude of EMG aftereffects (*p* = 0.24), to the adaptation (speed vs. *β*_adapt_, *p* = 0.81, speed vs. *β*_no-adapt_, *p* = 0.61), or to the magnitude of corrective responses upon ‘off’ transition 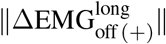 (*p* = 0.64). Interestingly, age-dependency was not observed in the magnitude of step-length asymmetry aftereffects (*ρ* = –0.364, *p* = 0.182, Figure 5D), which are conventionally used to characterize sensorimotor recalibration in locomotor. The partially distinct information characterized by muscle activity and step length asymmetry is further indicated by the poor relation between muscle activity aftereffects and kinematic aftereffects (*ρ* = 0.389, *p* = 0.152, not shown). In sum, we observed an age-related effect in the strucuture of corrective responses following a long exposure of the split-belts condition and smaller EMG aftereffects in older adults.

## 4 Discussion

We showed that muscle activity patterns following sudden changes in environmental walking conditions are adapted with prolonged exposure to a new environment. Importantly, the structure of corrective activity upon removing the novel split condition was similar to the muscle patterns expected in response to the introduction of a novel environment with dynamics opposite to the one originally experienced. We claim that these corrective responses post-adaptation can be interpreted as a proxy for sensorimotor recalibration since the adaptive structural features were only observed when subjects had enough time to adopt the split pattern as their new reference gait. Interestingly, older subjects showed less adaptation of this corrective activity, leading to smaller aftereffects in muscle space. Our results are relevant not only because we identified a sensitive measure of individual sensorimotor adaptation, but also because we provided a valuable characterization of normative changes in muscle activity from split-belts walking, which has been suggested as a rehabilitation therapy.

### 4.1 Learning of a new steady-state locomotor pattern

Muscle activity modulation during late split-belts walking was mostly anti-symmetric, likely due to the uneven demands of the split environment, such as the distinct speed-specific propulsion force demands for each leg (Sombric et al. 2018). However, motor patterns were not simply adjusted based on ipsilateral speed demands, but activity of proximal muscles seemed influenced by the speed of the other leg’s belt. This highlighted the bilateral nature of locomotion (Maclellan et al. 2014) and contrasted with the idea that each leg is adapted independently, as observed in hybrid split-belts walking (i.e., one leg moving forward fast and the other leg moving backwards slowly, Choi and Bastian 2007) possibly because of the peculiar demands of this hybrid task. In fact, most muscles increased activity when the split condition was introduced and plateaued at reduced activation levels, supporting the notion that initial stability demands result in higher activation levels that decreases as motor patterns become more efficient (Franklin et al. 2008, Huang and Ahmed 2014, Finley et al. 2013). Notably, the learned (steady-state) modulation pattern did not resemble the aftereffects in muscle activity, except in a few muscles. This was to a certain degree unexpected given that kinematic aftereffects are thought to reflect a continuation of the adapted motor commands in the altered environment (Malone et al. 2012, Morton and Bastian 2006, Ogawa et al. 2014). Plantarflexors (calf muscles) were the predominant muscles exhibiting similar activity before and after split-belts walking (only during double support). Plantarflexor activity contributes to displacing the body and leg forward (Neptune et al. 2001), and consequently modulate step length (Neptune et al. 2008). Thus, continuation of plantarflexors’ anti-asymmetric activity post-adaptation might lead to the known aftereffect on step length asymmetry following split-belts walking. To test this hypothesis, future work is needed with a different approach, such as musculoskeletal simulations (Steele et al. 2010, Song and Geyer 2015), which will enable investigating the EMG-kinematic relation during and after split-belts walking. In sum, muscle activity in the split condition was anti-symmetric and only plantarflexors’ modulation during late split-belts walking continued upon removal of the split environment.

### 4.2 Corrective activity post-adaptation indicates the motor system’s adapted state

We demonstrated that the structure of corrective activity upon removal of the novel split-belts environment indicated subjects’ adapted state and not just environmental changes. This was shown by the difference in the structure of corrective activity after identical environmental transitions following different exposure durations to the split environment. Corrective muscle activity following a long, but not a short, exposure period was equivalent to the modulation expected upon the introduction of an environment opposite to the one subjects experienced. This is consistent with the observation that removal of an altered environment facilitates meta-learning of an opposite environment (Herzfeld et al. 2014b). Our results suggest that subjects have adopted the gait pattern of late split-belts walking as a “new normal” such that deviations from it are experience as perturbations. This interpretation is supported by studies indicating that subjects shift their perception of what constitutes symmetric walking (i.e., same speed for two legs) following adaptation to the split-belts environment (Vazquez et al. 2015, Statton et al. 2018). It is worth pointing out that the observed structure of corrective responses was not exactly what we expected under our adaptive hypothesis: both *β*_adapt_ was smaller than 1, and *β*_no-adapt_ was larger than 0. We interpret the non-zero *β*_no-adapt_ as some amount of environment-based switching in corrective responses due to rapid strategic adjustments once subjects realize that they are no longer in the split condition. This is similar to re-aiming reaching direction when subjects notice that visuomotor rotations are removed (Morehead et al. 2015, McDougle et al. 2016). Moreover, the differences between the estimated and expected regressor values might be simply caused by methodological limitations. First, consider that the estimation of *β*_adapt_ values relies on our assumption that ΔEMG_on(–)_ can be inferred from ΔEMG_on(+)_. Thus, *β*_adapt_ (but not *β*_no-adapt_) can be biased downwards by any asymmetries between the legs due to mismatched placement of the EMG sensors or temporally misaligned activity originated by unequal half-cycle durations across the legs. Second, any noise in the regression factors (i.e., ΔEMG_on(+)_ and ΔEMG_on (–)_) biases the estimates of the regressor coefficients towards 0. This is because noise in a high-dimensional space is more likely to lead to misalignment between these regression vectors and the vector characterizing corrective activity post-adaptation 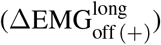. Both of these problems are more acute when estimating *β*_adapt_ and *β*_no-adapt_ per subject, which have larger noise levels and where activity is more likely to be asymmetrically recorded. Consistently, the effect of noise can be reduced by using a larger number of strides to characterize muscle activity during ‘early’ epochs. For example, similar results were obtained when using 15 strides (rather than 5) to compute corrective activity, but with slightly larger regression coefficients and *R*^2^. Despite these methodological limitations, we concluded that the structure of corrective activity post-adaptation reflects the recalibration of the motor system during split-belts walking because overall corrective responses in the ‘off’ transition were substantially more similar to the adaptive (O2) than environment-dependent (O1) corrective activity. Taken together our results suggests that the structure of corrective responses indicates the extent of sensorimotor recalibration at an individual level.

### 4.3 The adaptation of corrective responses suggests learning-dependent tuning of feedback activity

We believe that our results are evidence of the influence of predictive mechanisms on reactive control. We could not use conventional approaches to experimentally discern the adaptation of internal models for predictive (feedforward) control from online feedback corrections (Yousif and Diedrichsen 2012, Wagner and Smith 2008, Crevecoeur and Scott 2013, Cluff and Scott 2013) given the continuous nature of the walking task. We could only dissociate adjustments that are immediate from those that take multiple steps and presumably result from adaptation of internal models of the environment and/or body Roemmich and Bastian (2015), Reisman et al. (2009), Torres-Oviedo and Bastian (2010). We concede that the corrective responses that we characterize include a variety of mechanisms that have different neural substrates, such as monosynaptic reflexes, longer loop responses, and voluntary movement corrections. However, we interpret the reported structural changes of these corrective responses as learning-dependent adjustments of feedback processes involving supraspinal mechanisms (i.e., longer loop responses). This interpretation is in accordance with previous reaching studies suggesting shared internal models between feedforward and feedback motor control (Wagner and Smith 2008, Crevecoeur and Scott 2013, Cluff and Scott 2013). We discounted the possibility that split-belts walking modifies short-latency monosynaptic stretch reflexes since these can only change with extensive training (Thompson et al. 2009). Also, we reasoned that we are capturing little strategic adjustments (i.e. change in feedforward actions), such as goal-directed changes of foot landings (Matthis et al. 2017) or muscle activations passively driven by the environment. Namely, these kinds of modulation would predict more environment-based switching at the ‘off’ (long) transition and we only observe little of that (as quantified by the small *β*_no-adapt_ values). Taken together, our results provide further evidence that feedback control is influenced by the recalibration of processes involved in predictive motor control.

### 4.4 Age-related decline in the adaptive nature of feedback activity

Aging affects the adaptation of muscle activity after, but not during, split-belts walking, suggesting a possible dissociation between the update of feedback and feedforward activity. Motor learning was affected by age as indicated by the reduced magnitude of aftereffects and less mirroring in feedback activity postadaptation. It has been suggested that older subjects have noisier sensory information (Konczak et al. 2012), and thus, rely less on sensory input and more on predictive mechanisms (Wolpe et al. 2016). This would explain both the weaker feedback responses upon environmental transitions and reduced recalibration of internal models (i.e., smaller β_adapt_) in older subjects. Despite the age-related modulation of aftereffects in muscle activity, this age dependency is not observed in kinematic aftereffects (Sombric et al. 2017, Bruijn et al. 2012). EMG signals may be more sensitive to the recalibration of internal models because they are a closer correlate of neural activity than kinematics. Further, the magnitude of muscle patterns during late Long Exposure, which we presume to be mostly feedforward-generated activity, was not affected with age. This dichotomy in age effects on feedback vs. feedforward activity supports the partial dissociation of the adaptation of these two processes (Yousif and Diedrichsen 2012). We conclude that age-related sensory decline might contribute to motor learning deficits in older adults, which is observed in the adaptation of corrective responses, but not in steady state motor patterns.

### 4.5 Study implications

Our findings suggests that corrective responses could be used as an alternate approach for quantifying sensorimotor recalibration. In discrete behaviors such as eye or arm movements, aftereffects are typically quantified by the initial error (e.g. the initial direction of a thrown dart after adaptation) – since it reflects the newly acquired predictive model when the altered environment is removed. This is not easily quantifiable in walking because of its continuous nature, and because there is no explicit error measure. Our novel approach highlights that the recalibration of the predictive model can be quantified not only by the standard “aftereffect”, but by the subsequent corrective response elicited by an inadequate initial action. This corrective response is present in other motor adaptation tasks (e.g. reaching correction after starting with a wrong movement direction) where it may also be used as an indicator of the motor system’s implicit recalibration.

From a clinical perspective, split-belts walking can potentially correct gait asymmetry post-stroke (Reisman et al. 2013), but little is known about the changes in muscle activity that can be expected from this task. Unlike previous work (Dietz et al. 1994, Raja et al. 2013, Ogawa et al. 2014) we provide a systematic characterization of healthy motor patterns during distinct phases of the gait cycle throughout the adaptation and deadaptation process. These normative data allow for the assessment of adaptation deficits post-stroke and early evaluation of the therapeutic limits of split-belts walking. Importantly, we show that inter-subject learning capacities may be better captured through EMG-based measurements than conventional kinematic measurements. These measures suggest that learning might be limited in some patients simply because of their age. Lastly, our results indicate that aftereffects are strongly influenced by corrective responses to the sudden transition to tied-belts walking, rather than solely represent changes in feedforward circuits, which are arguably the true target of rehabilitation therapies. We conclude that the long-term rehabilitative potential of split-belts walking may not be well captured by solely measuring kinematic behavior immediately after adaptation.

## Acknowledgements

We thank Reza Shadmehr, Adrian Haith, and Ryan Roemmich for their comments on earlier versions of this manuscript. We thank the anonymous reviewers for their constructive criticism.

---

**Figure 1-1:**
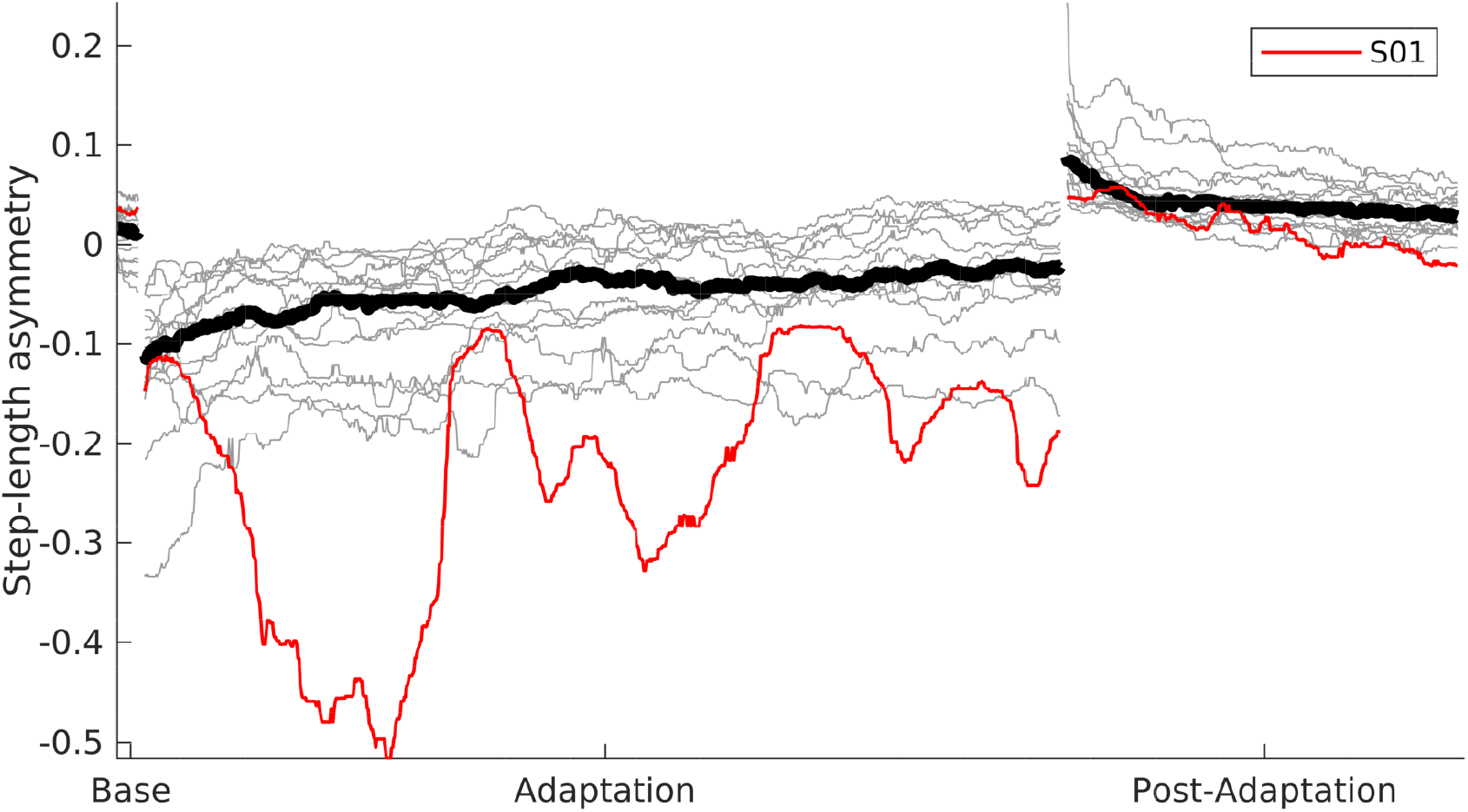
S01 is an outlier and failed to walk normally during Long Exposure. **(A)** Step-length asymmetry through Baseline, Long Exposure and Washout. Data was filtered with a running median filter of width 31 samples. S01 shown in red, other subjects in gray. Group median shown in black.

## Notes

**Conflict of Interest:** The authors declare no competing financial interests.

